# The long-term effects of genomic selection: 2. Changes in allele frequencies of causal loci and new mutations

**DOI:** 10.1101/2023.02.20.529287

**Authors:** Yvonne C.J. Wientjes, Piter Bijma, Joost van den Heuvel, Bas J. Zwaan, Zulma G. Vitezica, Mario P.L. Calus

**Affiliations:** Wageningen University & Research, Animal Breeding and Genomics, 6700 AH Wageningen, The Netherlands; Wageningen University & Research, Laboratory of Genetics, 6700 AH Wageningen, The Netherlands; INRAE, GenPhySE, 31326 Castanet-Tolosan, France

**Keywords:** Selection, Genomic changes, Allelic architecture, Mutations, Allele fixation, Non-additive effects, Genetic variance, Genomic selection

## Abstract

Genomic selection has become the dominant tool for genetic improvement in livestock and plants. Therefore, its sustainability is essential for global food production. Selection changes the allelic architecture of traits to create genetic gain. It remains unknown whether the changes in allele architecture are different for genomic selection and whether they depend on the genetic architectures of traits. Here we investigate the allele frequency changes of loci and new causal mutations under fifty generations of phenotypic, pedigree, and genomic selection, for a trait controlled by either additive, additive and dominance, or additive, dominance and epistatic effects. Genomic selection resulted in slightly larger and faster changes in allele frequencies of causal loci than pedigree selection. For each locus, allele frequency change per generation was not only influenced by its statistical additive effect, but also by the linkage phase with other loci and its allele frequency. Selection fixed a large number of loci, and five times more unfavorable alleles became fixed with genomic and pedigree selection than with phenotypic selection. For pedigree selection, this was mainly a result of increased genetic drift, while genetic hitchhiking had a large effect with genomic selection. When epistasis was present, the average allele frequency change was smaller (∼15% lower) and a lower number of loci became fixed for all selection methods. We conclude that for long-term genetic improvement, it is very important to be able to minimize the impact of hitchhiking and to limit the loss of favorable alleles more that current genomic selection methods do.

## INTRODUCTION

Genetic selection has been applied for many generations in animal and plant populations as well as in experimental species (*e.g.* mice, fruitflies, butterflies). This has resulted in a considerable improvement in desirable traits of those populations (Beniwal *et al*. 1992; Weber 1996; Dudley and Lambert 2003; Havenstein *et al*. 2003a, b). The aim of selection is to select the genetically most suitable individuals to produce the next generation. This will increase the frequency of alleles with a positive average effect on the desirable traits, which is known as genetic gain (Falconer and Mackay 1996; Walsh and Lynch 2018).

Selection has traditionally been based on pedigree and phenotypic information of the individuals and their relatives. In the last decade, pedigree selection has more and more been replaced by genomic selection (Meuwissen *et al*. 2001; Hayes *et al*. 2009; Knol *et al*. 2016; Meuwissen *et al*. 2016; Wolc *et al*. 2016). The use of genomic selection has likely accelerated the changes in allele frequencies across generations in certain regions on the genome (Heidaritabar *et al*. 2014; Liu *et al*. 2014; Doekes *et al*. 2018). It has been hypothesized that genomic selection mainly focusses on genes with a large contribution to the genetic variance (i.e., genes with a large effect and high minor allele frequency (MAF)) and tends to ignore the regions with a smaller contribution to the genetic variance of the trait of interest (Goddard 2009; Bijma 2012). As a consequence, the risk of losing rare favorable alleles has increased since the introduction of genomic selection (Jannink 2010; Liu *et al*. 2014; De Beukelaer *et al*. 2017). Monitoring and understanding the impact of selection on the changes in allele frequency of loci is important for investigating the long-term effects of selection. Since the vast majority of causal loci are unknown, simulation studies can contribute to better understand and quantify the changes at causal loci and how these are affected by selection approaches.

In a previous study (Wientjes *et al*. 2022), we have used simulations to investigate the long-term effects of selection on genetic gain, genetic variance and genetic architecture. The main conclusion was that the long-term response to selection was larger with phenotypic selection than with genomic selection, because genomic selection lost more genetic variance. Genomic selection always outperformed pedigree selection and lost a similar amount of genetic variance. Both genetic gain and loss of genetic variance depended on the presence of non-additive effects, such as dominance and epistasis. Especially when epistasis was present, selection changed the genetic architecture of the trait. The average allele frequency change was smaller with epistasis, because epistasis can change the direction of selection on an allele over time. However, we did not study the pattern of allele frequency change in detail in our previous study.

Although epistatic interactions between causal loci are known to be common (Carlborg and Haley 2004; Carlborg *et al*. 2006; Flint and Mackay 2009; Huang *et al*. 2012), not much is known about the complete genetic interaction network in animals and plants. More elaborate information is available for the genetic interaction network in yeast (Tong *et al*. 2004; Boone *et al*. 2007; Costanzo *et al*. 2016), where epistatic interactions were described between 90% of the identified causal loci. Most of the loci were involved in only a few interactions, while some loci were involved in many interactions. A similar pattern, although studied in less detail, is described for other laboratory species, such as *C. elegans* (Lehner *et al*. 2006), *Drosophila* (Huang *et al*. 2012), and mice (Tyler *et al*. 2017). Moreover, protein-protein binding interaction networks are similar in yeast, animals, including humans, and plants. Therefore, Boone *et al*. (2007) and Mackay (2014) argued that it is likely that the observed genetic interaction network in yeast may reflect such interaction networks in other species as well.

When investigating the long-term effects of selection, it is important to also consider new mutations. This is because after sufficient time, the response to selection will be completely driven by the variation due to new mutations (Walsh and Lynch 2018) and without mutations, the genetic variance will rapidly deplete. The potential of selection methods to maintain and utilize the variance generated by new mutations after 20 generations of selection differs between pedigree and genomic selection methods, and is especially limited when no own performance information is used in the selection criterion (Mulder *et al*. 2019). However, little is known about the potential of selection methods to maintain and exploit new mutations over longer time.

To date, the impact of the selection method and non-additive effects (dominance and epistasis) on the change in allelic architecture of complex traits under selection in the long-term are still unknown. More information about those changes is essential to better understand and predict the long-term impact of selection. Therefore, the aim of this study is to open up the black box of how selection affects the allelic architecture of complex traits. To this end, we simulated 50 generations of phenotypic, pedigree, and genomic selection for three genetic architectures with only additive, additive and dominance, or additive, dominance and epistatic effects and analysed the allelic architecture through the changes in allele frequencies of causal loci and new mutations.

## MATERIALS AND METHODS

We use the simulated data from our previous study (for details see (Wientjes *et al*. 2022)). In brief, a historical population was simulated in QMSim software (Sargolzaei and Schenkel 2009). From this historical population, a number of individuals was selected to form the starting population under selection, which was further simulated using our own developed Fortran program. The simulated data represents a livestock population in terms of allele frequency distribution, which was strongly U-shaped for segregating loci as typically observed in sequence data (Daetwyler *et al*. 2014; Eynard *et al*. 2015; Heidaritabar *et al*. 2016; Bolormaa *et al*. 2019). Moreover, the linkage disequilibrium (LD) pattern was comparable to livestock populations, with reasonably strong linkage at short distances on the genome (Andreescu *et al*. 2007; Badke *et al*. 2012; Veroneze *et al*. 2013) and (File S1; Figure S1.1 and Figure S1.2). In each generation, the best 100 females and 100 males were randomly mated using a mating ratio of 1:1 and a litter size of 10 (5 females and 5 males). In total, 15 scenarios were simulated that contained all combinations of three genetic models and five selection methods. All scenarios were replicated 20 times.

### Genome and mutations

The genome of the simulated population contained 10 chromosomes of 100 cM each, with on average one recombination event per offspring chromosome. Before selection, a marker set was selected of 20,000 segregating loci with a uniform allele frequency distribution. A set of 2,000 segregating causal loci was randomly selected with a U-shaped allele frequency distribution. Moreover, a set of 4000 non-segregating loci (i.e., that showed no variation at the start of selection) was randomly selected to serve as locations for causal mutations. Each individual had on average 0.6 new mutations, sampled from a Poisson distribution. This resulted in a mutational variance of 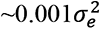 when assuming only additive effects (as explained later), which agrees with estimates from experimental populations (Hill 1982a; Houle *et al*. 1996; Lynch and Walsh 1998). For computational reasons, the loci and effects for new mutations were recycled while treating every time the mutation as a new mutation. Thus, for each new mutation, a locus that did not segregate at that moment was selected, while maximizing the time between two mutations at the same locus. This resulted in using each locus on average once in every 6-7 generations as a new mutation. We believe that the recycling of loci for mutations did not impact the results of our study, because the majority of mutations (80%) were already lost in the first generation as a result of drift and not related to the effect of the mutation (File S2; Table S2.1).

### Complex traits

For all causal loci (including loci for mutations), functional effects were assigned using one of three genetic models, namely a model with only additive effects (A), a model with additive and dominance effects (AD), and a model with additive, dominance and epistatic effects (ADE). Additive and dominance effects were simulated for all causal loci, based on established approaches (i.e., Wellmann and Bennewitz 2011; Duenk *et al*. 2020). Additive effects (*a*) were sampled from *N*(0,1). Dominance effects (*d*) were simulated to be proportional to the additive effects by first sampling a dominance degree (*dd*) from *N*(0.2,0.3) and subsequently using *d_i_* = *dd_i_*|*a_i_*| for all loci *i*.

Only pairwise epistatic effects were simulated based on the epistatic network described in yeast, where 90% of the causal loci were involved in one or more interactions (Costanzo *et al*. 2016). In this network, most loci were involved in a few interactions, and a few loci were involved in many interactions. Epistatic effects (*e*) were independently simulated for the nine possible genotype combinations of a pairwise interaction and scaled to be proportional to the additive effects of both loci. Thus, for the interaction between loci *i* and *j*, an epistatic degree (ε) was sampled from *N*(0, 0.45) for each possible genotype combination, and the epistatic effect was calculated as 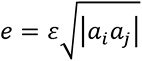.

Based on the functional effects and the genotypes, a total genotypic value was calculated for each individual. A residual, sampled from a normal distribution using a broad sense heritability of 0.4, was added to calculate the phenotypic value. The simulated functional additive, dominance, and epistatic effects were used to compute the statistical additive (α) and dominance (δ) effects for each causal loci based on the allele frequencies of the causal loci, using the Natural and Orthogonal Interaction Approach (NOIA) (Álvarez-Castro and Carlborg 2007; Vitezica *et al*. 2017). The statistical additive effects were used to compute the total additive genetic value (i.e., true breeding value) across all loci *i* of each individual as **A* =* ∑ *w_ai_α_i_*, with

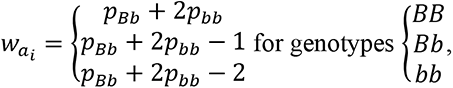

where *p_BB_*, *p_Bb_*, and *p_bb_* represent the frequencies of the genotypes *BB*, *Bb*, and *bb* for locus *B*. The additive genetic variance was the variance in *A* among individuals.

The statistical dominance effects were used to compute the total dominance deviation across all loci *i* of each individual as *D* = ∑ *w_di_δ_i_*, with

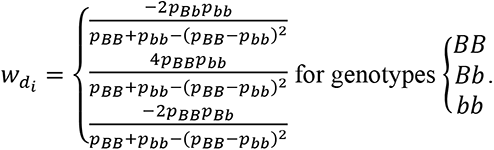

The dominance genetic variance was the variance in *D* among individuals. The total genetic variance minus the additive and dominance variance is the epistatic variance. For genetic model ADE, almost 50% of the variation at the functional level was a result of the epistatic effects, however, at the statistical level, more than 60% of the variance was additive and only 5% epistatic.

### Selection process

We used five artificial selection methods. The first method, RANDOM, selected in each generation randomly the parents of the next generation. The second method, MASS, selected the parents based on their own phenotypes. The other three methods selected parents based on estimated breeding values using Best Linear Unbiased Prediction (BLUP).

With PBLUP_OP, breeding values were estimated using pedigree information from the last eight generations and phenotypes from the last three generations, including own performances (OP) of the selection candidates. With GBLUP_NoOP and GBLUP_OP, breeding values were estimated using marker genotypes and phenotypes from the last three generations, either excluding (NoOP) or including (OP) the own performance of the selection candidates. Breeding values were estimated with the MTG2 software (Lee and van der Werf 2016), simultaneously with the variance components. Since commercial models for genetic improvement of animals and plants are mainly based on additive models (Crossa *et al*. 2010; Goddard *et al*. 2010), our breeding value estimation model included a fixed mean, random additive genetic effects, random litter effects and residuals. The random litter effect was included to prevent that the resemblance between full sibs due to non-additive genetic effects created bias in the estimated variance components and breeding values.

### Change in allelic architecture

We investigated the changes in allelic architecture of causal loci over 50 generations of drift and selection of a complex trait. First, we investigated the general pattern of change in allele frequency across the genome and the relation between allele frequency change and statistical additive effects (i.e., allele substitution effects, α). However, it is well known that linked loci can have a strong impact on the selection pressure on a locus and thereby also on allele frequency change and probability of fixation (Walsh and Lynch 2018). Therefore, we also investigated the relationship between allele frequency change and the *apparent* effect of a causal locus, which also included the effects of linked loci. Apparent effects were estimated in each generation for each locus as the *simple* regression coefficient of the true breeding value on the gene content at the locus, ***TBV*** = intercept + ***z***_i_*α*_Apparent,i_ + ***e***, where ***TBV*** is a vector with the true breeding values of all individuals in one generation, ***z****_i_* is a vector with the allele count of locus *i* of all individuals, α_Apparent,i_ is the apparent effect of locus *i*, and ***e*** is a vector with residuals. Note that the ordinary statistical additive effect (α) is the *partial* regression coefficient of the total genotypic value on gene content, ***TGV** = intercept + **z**_i_α_i_ + ∑_j≠i_ **z**_j_α_j_ + **e***.

Then, we zoomed in on the causal loci with either a large (>0.9) or small (<0.01) change in allele frequency across the 50 generations of selection and investigated the characteristics of those loci, such as average functional and statistical effects, interaction network, and allele frequency pattern. The cut-off values for the change in allele frequencies were chosen such that a reasonable number of loci met the criteria (See File S1; Figure S1.3 for the distribution in allele frequency change). The number of loci with an allele frequency change < 0.01 was very large, because loci initially close to fixation could not change in allele frequency after becoming fixed. Therefore, we used an additional criterion that the starting MAF had to be at least 0.05 for the loci with a small allele frequency change.

Thereafter, we investigated the characteristics of the causal loci that became fixed or lost during selection, while using the criterion that the minimal change in allele frequency had to be 0.2, to disregard loci that were already close to fixation or loss in the initial generation. In addition, we looked at the probability of loci to become fixed or lost over 50 generations of selection as a function of their starting allele frequency, and compared the pattern with the expected pattern under drift. The expected pattern under drift was obtained from additional simulations with no selection and the same effective population size (*N_e_*). The *N_e_* for each selection scenario was estimated from the pedigree kinship coefficient, using 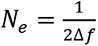 (Caballero 1994), where *Δ*f** is the rate of kinship based on the off-diagonal elements of the pedigree relationship matrix. Then new simulations were run with random selection and the same *N_e_* as in the selection scenarios, by adjusting the number of parents (File S3).

Finally, we investigated the number of mutations that appeared between generation 0 and 49, and still segregated in generation 50, as well as their allele frequency in generation 50.

### Data availability

File S4 contains the QMSim input file, Fortran programs and seeds used to select the markers and causal loci, to simulate functional effects and genotypes and phenotypic values of new generations, and the interaction matrix used to simulate epistatic effects.

## RESULTS

### General characteristics of selection

For ease of understanding the results, Table 1 shows the change in allele frequency and phenotypic value, and the proportion of genetic variance lost, with more details in (Wientjes *et al*. 2022). When only additive (model A) or additive and dominance (model AD) effects were present in the population, the overall phenotypic change over 50 generations of selection was highest for GBLUP_OP, followed by MASS, PBLUP_OP and GBLUP_NoOP (Table 1). When also epistatic effects were present (model ADE), MASS selection outperformed GBLUP_OP after ∼45 generations. As expected, RANDOM selection did not change the average phenotype over the 50 generations. The genetic variance was constant under RANDOM selection, while ∼60% of the genetic variance was lost with MASS selection. The other selection methods resulted in losing 70-85% of the genetic variance, and lost slightly more genetic variance when only additive effects were present.

**TABLE 1.** Change in average phenotype, genetic variance and allele frequency across 50 generations of selection for the five selection methods and three genetic models^1^. The five selection methods were: RANDOM selection, MASS selection, PBLUP selection with own performance (PBLUP_OP), GBLUP selection without own performance (GBLUP_NoOP) or with own performance (GBLUP_OP). The three genetic models were a model with only additive effects (A), with additive and dominance effects (AD), or with additive, dominance and epistatic effects (ADE).

|  | Phenotypic change<sup>2</sup> | Proportion of genetic variance lost | Average change in allele frequency |
| --- | --- | --- | --- |
| *Model A* |  |  |  |
| **RANDOM** | 0.02 (0.11) | 0.03 (0.03) | 0.023 (0.000) |
| **MASS** | 27.31 (0.44) | 0.60 (0.01) | 0.054 (0.001) |
| **PBLUP_OP** | 27.02 (0.46) | 0.79 (0.01) | 0.062 (0.000) |
| **GBLUP_NoOP** | 25.25 (0.45) | 0.82 (0.01) | 0.063 (0.001) |
| **GBLUP_OP** | 29.37 (0.42) | 0.83 (0.00) | 0.065 (0.001) |
| *Model AD* |  |  |  |
| **RANDOM** | 0.02 (0.12) | 0.02 (0.03) | 0.023 (0.000) |
| **MASS** | 26.80 (0.37) | 0.56 (0.02) | 0.053 (0.000) |
| **PBLUP_OP** | 25.43 (0.37) | 0.74 (0.01) | 0.061 (0.000) |
| **GBLUP_NoOP** | 23.54 (0.34) | 0.81 (0.01) | 0.064 (0.000) |
| **GBLUP_OP** | 27.68 (0.29) | 0.81 (0.01) | 0.065 (0.000) |
| *Model ADE* |  |  |  |
| **RANDOM** | 0.05 (0.11) | 0.00 (0.03) | 0.023 (0.000) |
| **MASS** | 20.87 (0.27) | 0.56 (0.02) | 0.042 (0.000) |
| **PBLUP_OP** | 18.76 (0.29) | 0.74 (0.01) | 0.053 (0.000) |
| **GBLUP_NoOP** | 16.79 (0.21) | 0.75 (0.01) | 0.054 (0.000) |
| **GBLUP_OP** | 20.21 (0.26) | 0.76 (0.01) | 0.055 (0.000) |

<sup>1</sup> Results are shown as averages across replicates with standard errors across replicates between brackets.
<sup>2</sup> Expressed in base generation additive genetic standard deviations.

### Change in allelic architecture

With RANDOM selection, the average absolute change in allele frequency across all causal loci was 0.023, which represents the effect of drift (Table 1). Selection increased the change in allele frequency. Under genetic models A and AD, the change in allele frequency was ∼2.3 times higher with MASS, ∼2.7 times higher with PBLUP_OP, ∼2.8 times higher with GBLUP_NoOP, and ∼2.9 times higher with GBLUP_OP than the change due to drift alone. The change in allele frequencies was substantially less (∼15% lower) when epistatic effects were present. Since MASS had the lowest average change in allele frequency and among the highest cumulative genetic gains (Table 1), MASS showed the smallest change in allele frequency per unit of genetic gain.

Figure 1 shows the absolute change in allele frequency for each locus. For RANDOM, most allele frequency changes were in the range of 0 to 0.25, with only very few changes above 0.4. With selection, allele frequency changes up to 1 were observed, indicating that some (0.02% for PBLUP_OP, GBLUP_NoOP and GBLUP_OP, 0.00% for MASS) new mutations became fixed within 50 generations. The vast majority (∼80%) of allele frequency changes were, however, still in the range of 0 to 0.25 with selection.

**FIGURE 1.**
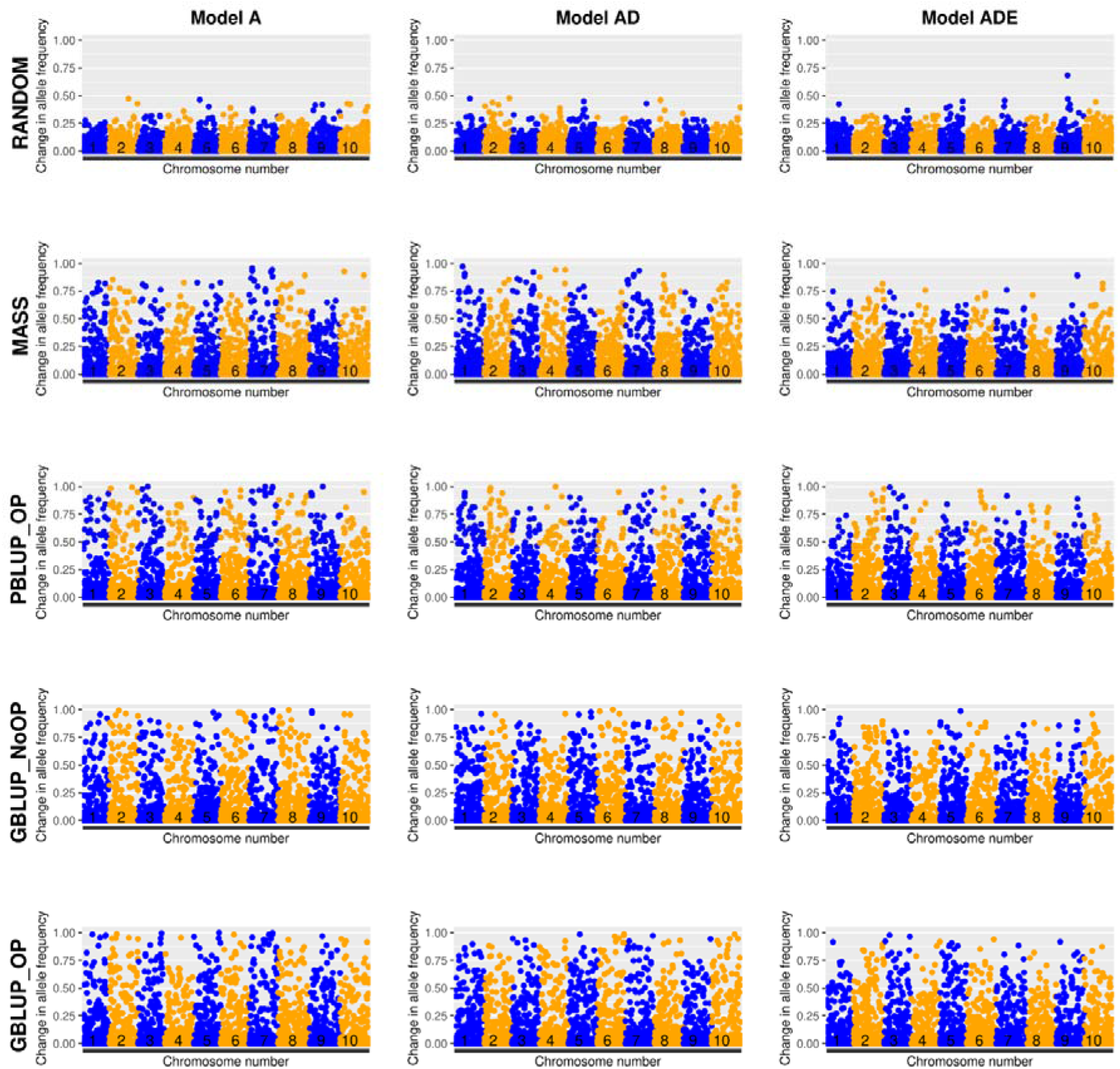
Absolute change in allele frequency of causal loci over 50 generations of selection versus the position on the genome for the five selection methods and three genetic models. The five selection methods were: RANDOM selection, MASS selection, PBLUP selection with own performance (PBLUP_OP), GBLUP selection without own performance (GBLUP_NoOP) or with own performance (GBLUP_OP). The three genetic models were a model with only additive effects (A), with additive and dominance effects (AD), or with additive, dominance and epistatic effects (ADE). Results are shown for one replicate.

The average change in allele frequency was larger when the initial heterozygosity at the locus was higher, and this relationship became stronger when selection was more accurate (File S1; Figure S1.4). With random selection, the maximum change in allele frequency was larger for loci with a higher initial heterozygosity. With selection, the maximum achieved change in allele frequency was closer to the maximum possible change in allele frequency and, therefore, higher for loci with a lower initial expected heterozygosity.

No clear peaks of adjacent causal loci that all had a large change in allele frequency were observed among causal loci (Figure 1). This is likely a result of the relatively low level of LD between causal loci (File S1; Figure S1.1). For the genomic selection scenarios, we also investigated the allele frequency changes of the markers, for which the level of LD was higher (File S1; Figure S1.2). By combining those with the allele frequency changes at causal loci, some peaks became visible where a causal locus and its surrounding markers together changed substantially in allele frequency (File S1; Figure S1.5).

### Allele frequency change *versus* effect size of loci

As expected, in the RANDOM scenario the average change in allele frequency, represented by the black dots in Figure 2, was around zero for all bins of causal loci based on their base-generation statistical additive effect. For the scenarios with selection, loci with a larger statistical additive effect showed on average a larger change in allele frequency. Moreover, the maximum negative change in allele frequency (i.e., an increase in frequency of the unfavorable allele) was smaller for loci with a larger statistical additive effect. The presence of epistatic effects made this trend less clear, as is also visible in the correlation coefficients between the change in allele frequency and the statistical additive effect in generation 0, which were lower for model ADE (File S2; Table S2.2). This observation is likely a result of changes in the statistical additive effect across generations which resulted in a large proportion of loci changing bin number (24.7 – 43.0% of loci after 10 generations, and 37.9 – 58.2% after 50 generations, File S2; Table S2.3).

**FIGURE 2.**
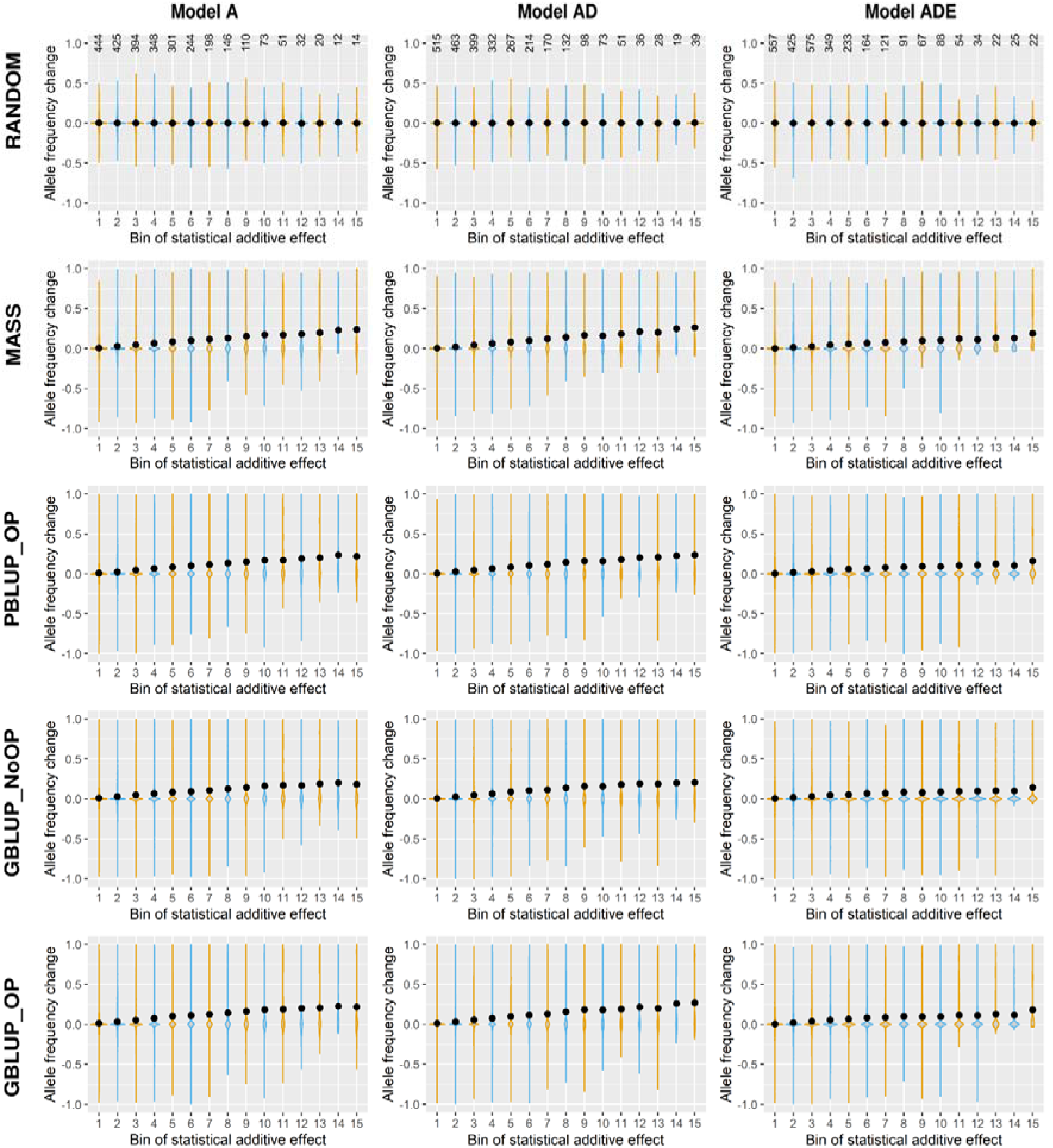
Change in allele frequency versus the size of the base-generation statistical additive effect (alpha), for the five selection methods and three genetic models. Loci are divided in bins based on their statistical additive effect in generation 0, where bin 1 contains the loci with the smallest statistical additive effect and bin 15 contains the loci with the largest statistical additive effect. Positive allele frequency changes indicate an increase in frequency of the favorable allele and negative allele frequency changes an increase in frequency of the unfavorable allele based on the allele substitution effects in generation 0. The five selection methods were: RANDOM selection, MASS selection, PBLUP selection with own performance (PBLUP_OP), GBLUP selection without own performance (GBLUP_NoOP) or with own performance (GBLUP_OP). The three genetic models were a model with only additive effects (A), with additive and dominance effects (AD), or with additive, dominance and epistatic effects (ADE). Results are given across all 20 replicates. The average number of loci per replicate in each bin is indicated at the top.

The regression coefficients of the change in allele frequency on the scaled allele substitution effect ranged from 1.5 to 2.2 for genetic models A and AD, and from 0.8 to 1.1 for genetic model ADE (File S2; Table S2.2). However, the correlation between the allele frequency change from one generation to the next and the statistical additive effect was very low for all selection methods in all generations (<0.05, File S1; Figure S1.6).

Loci with larger statistical additive effects on average also had larger apparent effects (File S5). However, the correlation between the statistical additive effect and the apparent effects within a generation was always below 0.2 (Figure 3). These results indicate that the apparent effect of a locus is for a larger part determined by the linkage with other loci than by the statistical additive effect. This linkage is partly a result of the LD pattern, but might also be a result of sampling in a small population. When only including loci with a MAF above 0.05, the correlation between the statistical additive effects and apparent effects became stronger (∼0.3 to ∼0.4, File S1; Figure S1.7). This is because the probability of all individuals that contain a specific allele to have an above or below average breeding value by chance becomes larger when the allele frequency is lower, which is reflected in the apparent effect. So, the impact of other loci on the apparent effect of a locus is larger when a locus has a lower MAF.

**FIGURE 3.**
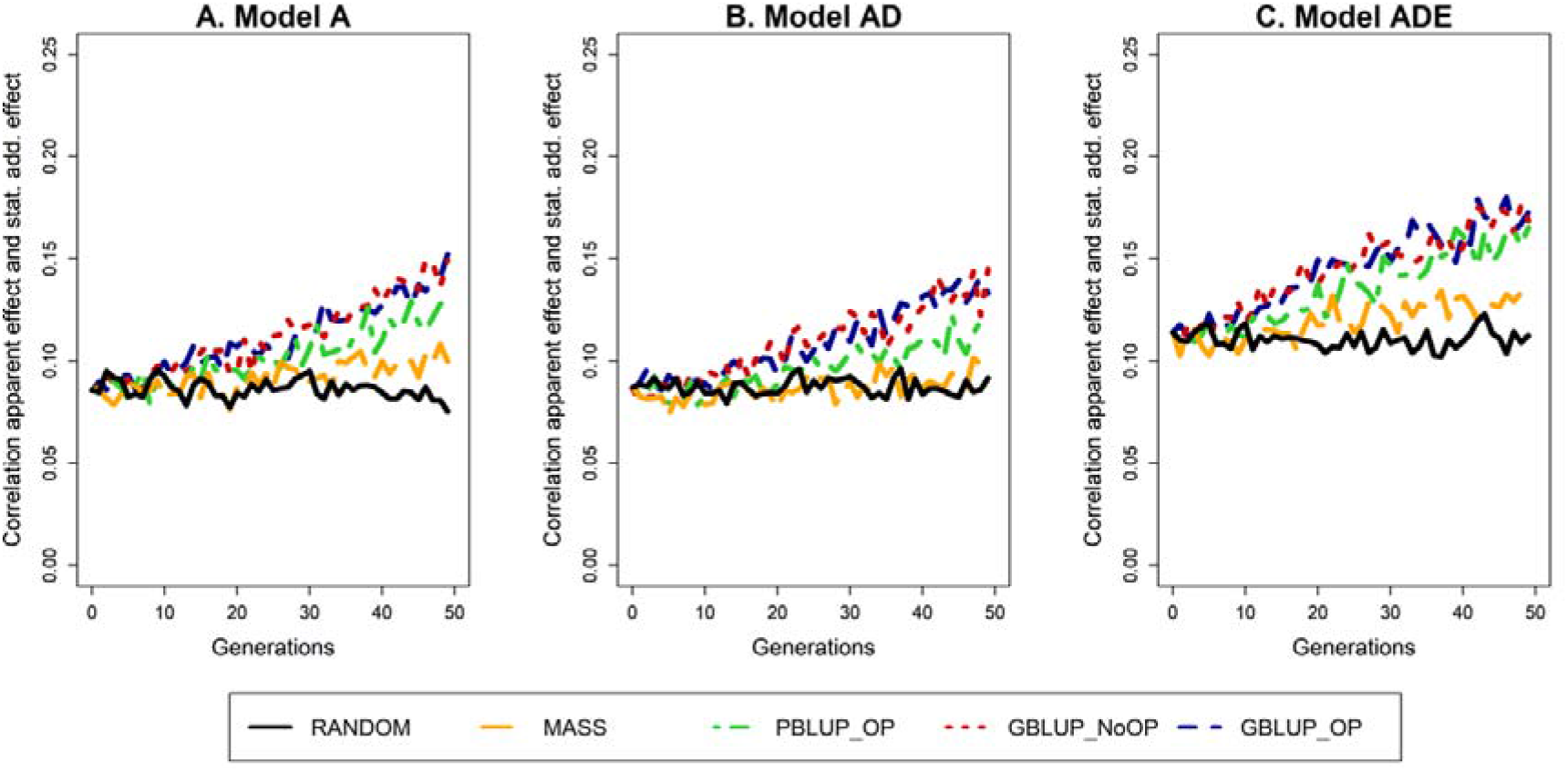
Correlation between the apparent effect and the statistical additive effect across generations for the five selection methods and three genetic models. The five selection methods were: RANDOM selection, MASS selection, PBLUP selection with own performance (PBLUP_OP), GBLUP selection without own performance (GBLUP_NoOP) or with own performance (GBLUP_OP). The three genetic models were a model with only additive effects (A), with additive and dominance effects (AD), or with additive, dominance and epistatic effects (ADE). Results are shown as averages of 20 replicates.

In the first generations of selection, the correlation between the apparent effect and statistical additive effect was similar across selection methods and genetic models underlying the trait (File S1; Figure S1.7). After more than ∼30 generations, the correlation seems to become slightly larger for the GBLUP and PBLUP models, especially for the additive model. This is probably a result of a reduction in the number of segregating causal loci as a result of selection, which increases the average distance between neighboring causal loci, thereby reducing the impact of LD.

The apparent effect of an allele correlated much stronger with allele frequency change than the statistical additive effect (Figure 4). The correlation between allele frequency change and apparent effect was highest for GBLUP_OP and lowest for MASS in the first generations of selection. Over generations, this correlation decreased, which happened at a faster rate for the GBLUP methods than for MASS and PBLUP. Moreover, the correlation was lower when non-additive effects, especially epistasis, was present.

**FIGURE 4.**
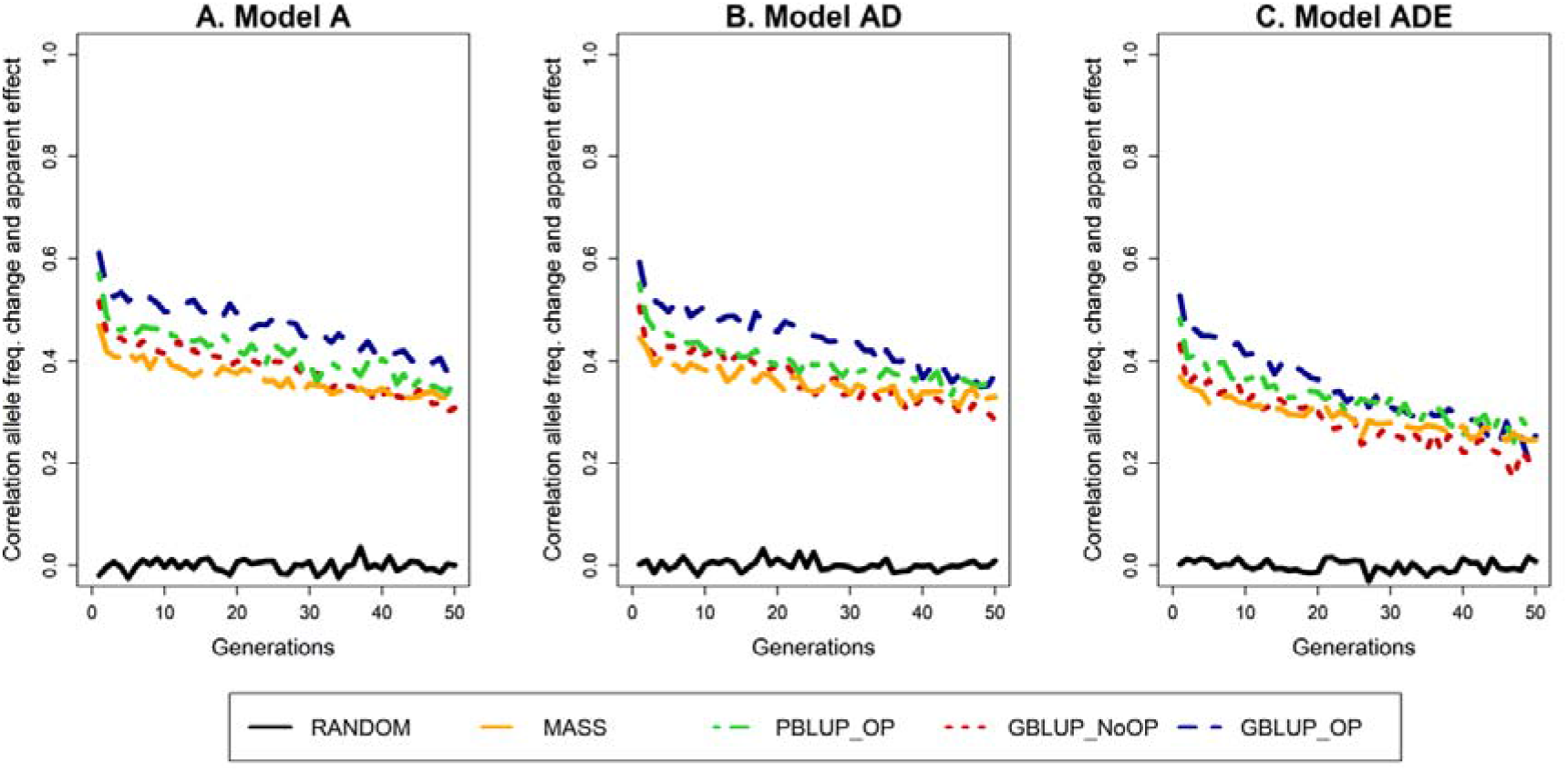
Correlation between the change in allele frequency towards the next generations and the apparent effect for the five selection methods and three genetic models. The change in allele frequency is expressed as the absolute change in allele frequency from generation *i* to generation *i* +1 divided by *p_i_*(1-*p_i_*), where *p_i_* is the allele frequency in generation *i*. The five selection methods were: RANDOM selection, MASS selection, PBLUP selection with own performance (PBLUP_OP), GBLUP selection without own performance (GBLUP_NoOP) or with own performance (GBLUP_OP). The three genetic models were a model with only additive effects (A), with additive and dominance effects (AD), or with additive, dominance and epistatic effects (ADE). Results are shown as averages of 20 replicates.

### Characteristics of loci with a large or small change in allele frequency

For RANDOM, there were no causal loci with a large (>0.9) change in allele frequency (Figure 5, Table 2, File S2; Table S2.4). With MASS, only few causal loci (≤0.1%; ≤6 out of 6000 loci) had a large change in allele frequency. With the other selection methods, ∼0.3% (17 out of 6000) to ∼0.5% (31 out of 6000) of the causal loci showed a large change in allele frequency under both model A and AD, and ∼0.1% (8 out of 6000) under model ADE. Across all genetic models, the GBLUP methods resulted in ∼24% more loci with a large change in allele frequency than PBLUP_OP. With MASS selection, almost all loci with a large change in allele frequency were changed in the favorable direction (∼99%). This was not the case for the other selection methods, where the frequency of the unfavorable allele increased by more than 0.9 for 2-19% of the loci, especially when epistasis was present.

**TABLE 2.** Characteristics of loci with a large (>0.9) or small (<0.01) change in allele frequency across 50 generations of selection for the five tion methods and three genetic models^1^. The five selection methods were: RANDOM selection, MASS selection, PBLUP selection with performance (PBLUP_OP), GBLUP selection without own performance (GBLUP_NoOP) or with own performance (GBLUP_OP). The genetic models were a model with only additive effects (A), with additive and dominance effects (AD), or with additive, dominance and atic effects (ADE).

|  | All loci |  |  |  | Loci with large (>0.9) change in allele frequency |  |  |  |  | Loci with small (<0.01) change in allele frequency |  |  |  |
| --- | --- | --- | --- | --- | --- | --- | --- | --- | --- | --- | --- | --- | --- |
| | Total Nr.<sup>2</sup> | Avg. add. effect<sup>3</sup> | Avg. Dom. effect<sup>4</sup> | Avg. $\alpha/\sigma_G$ in gen. 0<sup>5</sup> | Total Nr.<sup>2</sup> | Prop. increase favorable allele | Avg. add. effect<sup>3</sup> | Avg. dom. effect<sup>4</sup> | Avg. $\alpha/\sigma_G$ in gen. 0<sup>5</sup> | Total Nr.<sup>2</sup> | Avg. add. effect<sup>3</sup> | Avg. dom. effect<sup>4</sup> | Avg. $\alpha/\sigma_G$ in gen. 0<sup>5</sup> |
| *Model A* |  |  |  |  |  |  |  |  |  |  |  |  |  |
| **RANDOM** | 6000 | 0.80 | - | 0.041 | 0 | - | - | - | - | 63 | 0.83 | - | 0.042 |
| **MASS** | 6000 | 0.80 | - | 0.041 | 6 | 0.99 | 1.92 | - | 0.098 | 18 | 0.56 | - | 0.029 |
| **PBLUP_OP** | 6000 | 0.80 | - | 0.041 | 24 | 0.98 | 1.52 | - | 0.077 | 7 | 0.66 | - | 0.033 |
| **GBLUP_NoOP** | 6000 | 0.80 | - | 0.041 | 25 | 0.93 | 1.30 | - | 0.066 | 7 | 0.58 | - | 0.029 |
| **GBLUP_OP** | 6000 | 0.80 | - | 0.041 | 31 | 0.96 | 1.47 | - | 0.075 | 7 | 0.51 | - | 0.026 |
| *Model AD* |  |  |  |  |  |  |  |  |  |  |  |  |  |
| **RANDOM** | 6000 | 0.80 | 0.23 | 0.039 | 0 | - | - | - | - | 65 | 0.79 | 0.24 | 0.038 |
| **MASS** | 6000 | 0.80 | 0.23 | 0.039 | 4 | 1.00 | 1.79 | 0.49 | 0.102 | 21 | 0.57 | 0.18 | 0.027 |
| **PBLUP_OP** | 6000 | 0.80 | 0.23 | 0.039 | 17 | 0.97 | 1.45 | 0.43 | 0.085 | 10 | 0.65 | 0.20 | 0.032 |
| **GBLUP_NoOP** | 6000 | 0.80 | 0.23 | 0.039 | 23 | 0.92 | 1.27 | 0.37 | 0.072 | 7 | 0.66 | 0.20 | 0.032 |
| **GBLUP_OP** | 6000 | 0.80 | 0.23 | 0.039 | 26 | 0.96 | 1.46 | 0.43 | 0.084 | 7 | 0.58 | 0.19 | 0.028 |
| *Model ADE* |  |  |  |  |  |  |  |  |  |  |  |  |  |
| **RANDOM** | 6000 | 0.80 | 0.23 | 0.034 | 0 | - | - | - |  | 65 | 0.80 | 0.23 | 0.027 |
| **MASS** | 6000 | 0.80 | 0.23 | 0.034 | 1 | 0.97 | 1.58 | 0.38 | 0.100 | 29 | 0.76 | 0.22 | 0.017 |
| **PBLUP_OP** | 6000 | 0.80 | 0.23 | 0.034 | 8 | 0.85 | 1.15 | 0.32 | 0.090 | 11 | 0.78 | 0.23 | 0.018 |
| **GBLUP_NoOP** | 6000 | 0.80 | 0.23 | 0.034 | 8 | 0.81 | 1.11 | 0.32 | 0.075 | 11 | 0.73 | 0.23 | 0.021 |
| **GBLUP_OP** | 6000 | 0.80 | 0.23 | 0.034 | 9 | 0.81 | 1.16 | 0.32 | 0.087 | 11 | 0.81 | 0.23 | 0.017 |

<sup>1</sup> Results are shown as averages across replicates. For standard errors of the mean, see File S2; Table S2.4.
<sup>2</sup> Total number of loci.
<sup>3</sup> Average absolute functional additive effect of loci that became fixed in the population.
<sup>4</sup> Average absolute functional dominance effect of loci that became fixed in the population.
<sup>5</sup> Average absolute statistical additive effect ( $\alpha$ ) divided by the total genetic standard deviation in generation 0 of loci that became fixed in the population.

**FIGURE 5.**
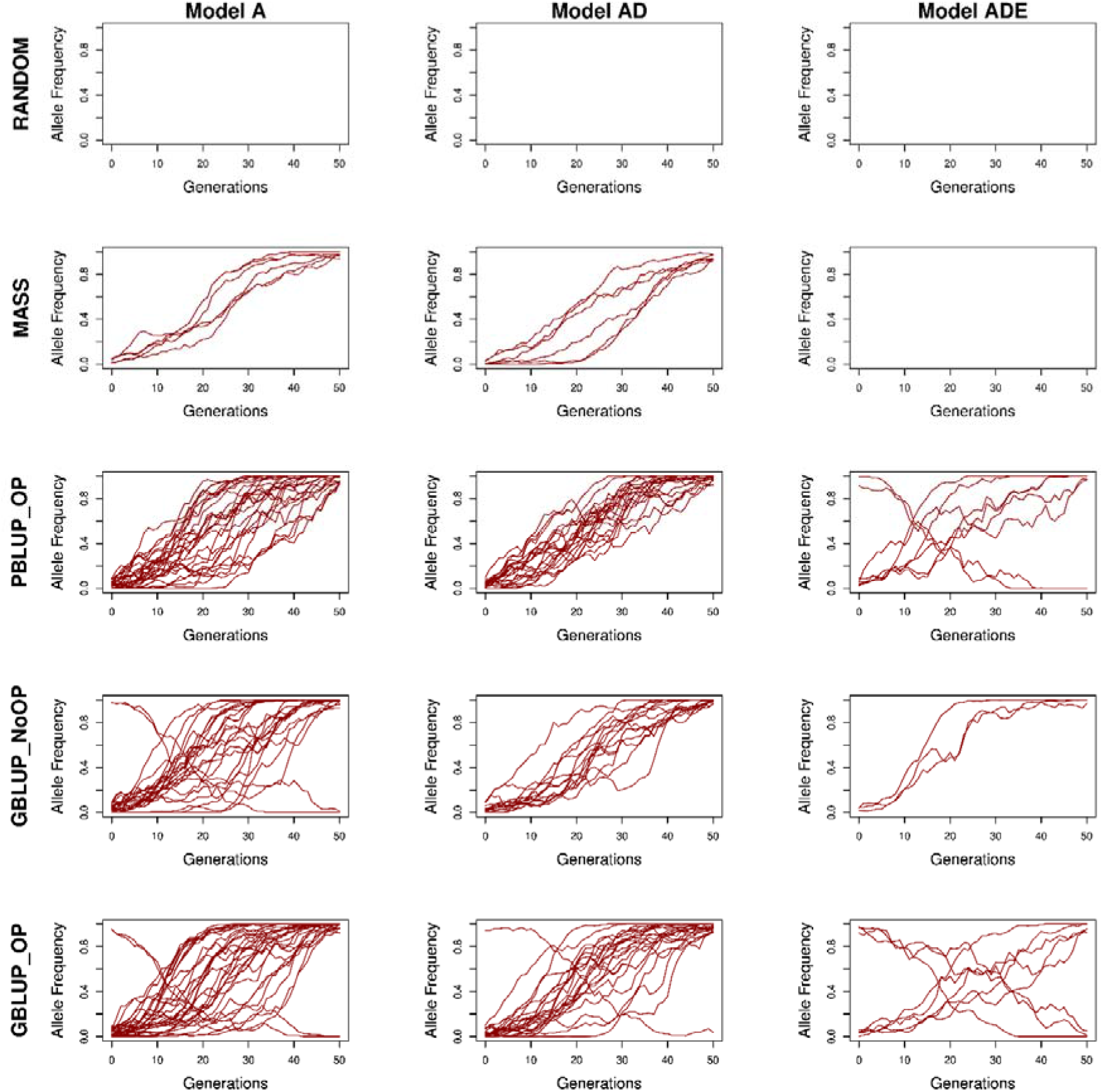
Trend in allele frequency of loci with a large change in allele frequency (>0.9) over the 50 generations of selection for the five selection methods and three genetic models. For all loci, the positive allele was counted, based on the statistical additive effect in generation 0. The five selection methods were: RANDOM selection, MASS selection, PBLUP selection with own performance (PBLUP_OP), GBLUP selection without own performance (GBLUP_NoOP) or with own performance (GBLUP_OP). The three genetic models were a model with only additive effects (A), with additive and dominance effects (AD), or with additive, dominance and epistatic effects (ADE). Results are shown for one replicate.

On average, the loci with a large change in allele frequency had a larger functional additive and dominance effect than an average locus (Table 2). Moreover, those loci had a larger statistical additive effect in generation 0. This was most pronounced with MASS, followed by PBLUP_OP and GBLUP_OP, and least pronounced with GBLUP_NoOP. Moreover, the loci with a large change in allele frequency were on average more involved in interactions compared to an average locus (Table 3).

**TABLE 3.** Average number of interactions per locus for all loci, and loci with a large (>0.9) or small (<0.01) change in allele frequency across 50 generations of selection for the five selection methods and the genetic model with additive, dominance and epistatic effect^1^. The five selection methods were: RANDOM selection, MASS selection, PBLUP selection with own performance (PBLUP_OP), GBLUP selection without own performance (GBLUP_NoOP) or with own performance (GBLUP_OP).

|  | **Average number of interactions of loci** |  |  |
| --- | --- | --- | --- |
|  | **All loci** | **Loci with large (>0.9) change in allele frequency** | **Loci with small (<0.01) change in allele frequency** |
| **RANDOM** | 121 (0.4) | - - | 85 (7.3) |
| **MASS** | 121 (0.4) | 459 (60.1) | 105 (5.4) |
| **PBLUP_OP** | 121 (0.4) | 234 (20.1) | 85 (7.3) |
| **GBLUP_NoOP** | 121 (0.4) | 193 (16.2) | 113 (9.9) |
| **GBLUP_OP** | 121 (0.4) | 243 (18.5) | 93 (8.5) |

<sup>1</sup> Results are shown as averages across the loci over all replicates with their corresponding standard errors of the mean between brackets.

Only few loci (0.1 – 1%) showed a small change in allele frequency (<0.01) over 50 generations (Table 2). Even though the change in allele frequency from generation 0 to 50 was small, allele frequencies fluctuated considerably (File S1; Figure S1.8). This result indicates that an allele frequency change <0.01 is largely due to chance, which agrees with the relatively small functional dominance effects for those loci (Table 2), indicating a very limited role for overdominance as a mechanism preventing change in allele frequency.

### Causal loci becoming fixed

With RANDOM, only ∼0.03% (2 out of 6000) of the loci changed at least 0.2 in allele frequency and became fixed over 50 generations (Figure 6; Table 4). This was much higher with MASS (2.2%; 133 out of 6000, model A), and even higher with PBLUP_OP, GBLUP_NoOP and GBLUP_OP (∼6%; 357 out of 6000, model A). When non-additive effects were present, the number of loci that became fixed was substantially lower (50% lower with MASS and 30% lower with PBLUP_OP, GBLUP_NoOP and GBLUP_OP for model ADE compared to model A). With MASS, the loci that were fixed were more frequently fixed for the favorable allele (82-89% of the cases) compared to the other selection methods (∼80% under model A and AD, and ∼70% under model ADE). For all selection methods, the loci fixed for the favorable allele had an above-average statistical additive effect (Tables 2 and 4). The average starting allele frequency of the favorable alleles that became fixed was highest for RANDOM (∼0.75), followed by MASS (∼0.61), and lowest for PBLUP_OP, GBLUP_NoOP and GBLUP_OP (∼0.54), indicating that the average change in allele frequency at loci where an allele became fixed was largest for PBLUP_OP, GBLUP_NoOP and GBLUP_OP.

**FIGURE 6.**
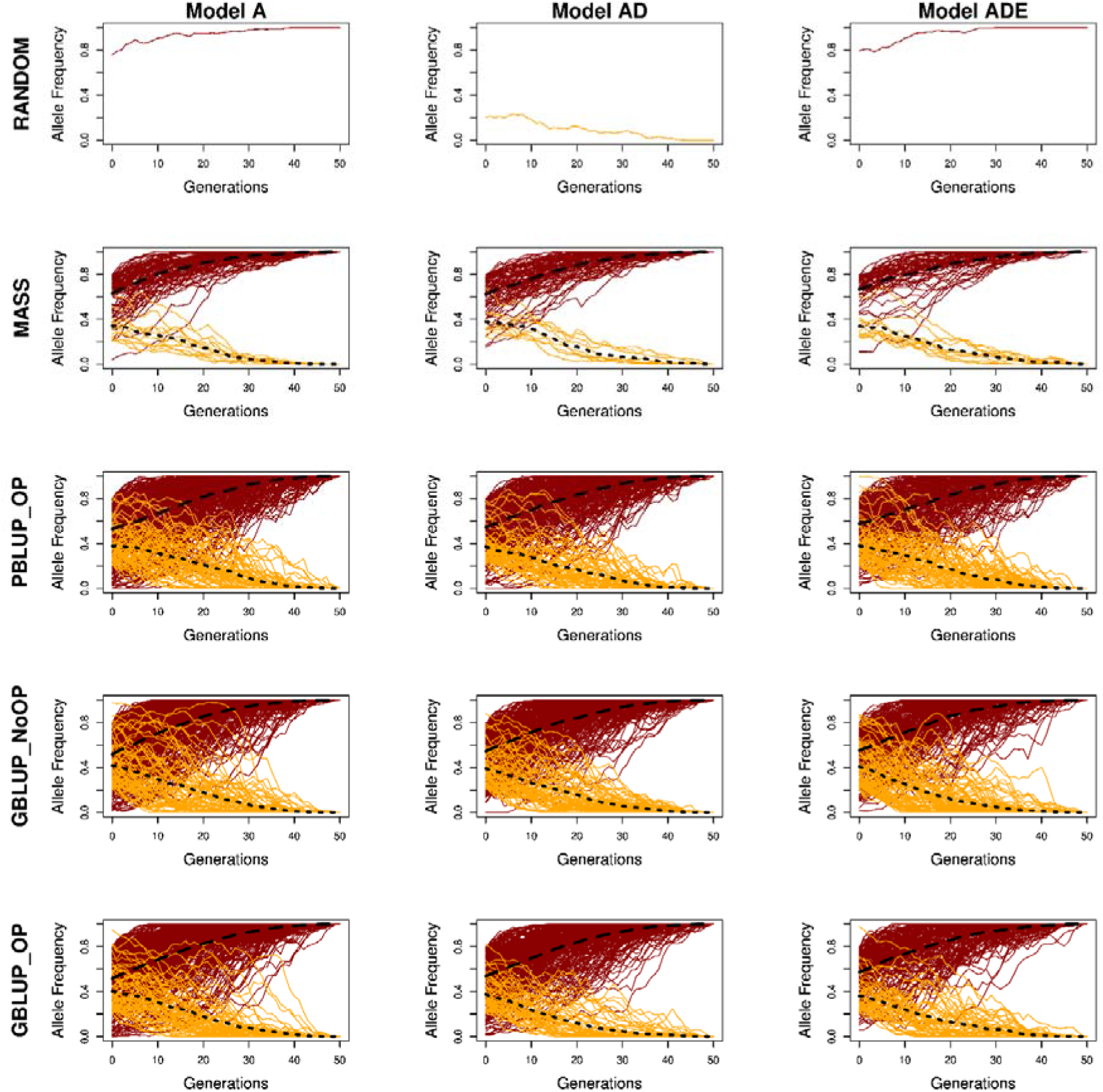
Trend in allele frequency of loci that become fixed in the 50 generations of selection for the five selection methods and three genetic models. For all loci, the favorable allele was counted, therefore red lines indicate loci fixed for the favorable allele and yellow lines indicate loci fixed for the unfavorable allele, based on the statistical additive effect in generation 0. The minimum change in allele frequency was set at 0.2. The five selection methods were: RANDOM selection, MASS selection, PBLUP selection with own performance (PBLUP_OP), GBLUP selection without own performance (GBLUP_NoOP) or with own performance (GBLUP_OP). The three genetic models were a model with only additive effects (A), with additive and dominance effects (AD), or with additive, dominance and epistatic effects (ADE). Results are shown for one replicate.

**TABLE 4.**
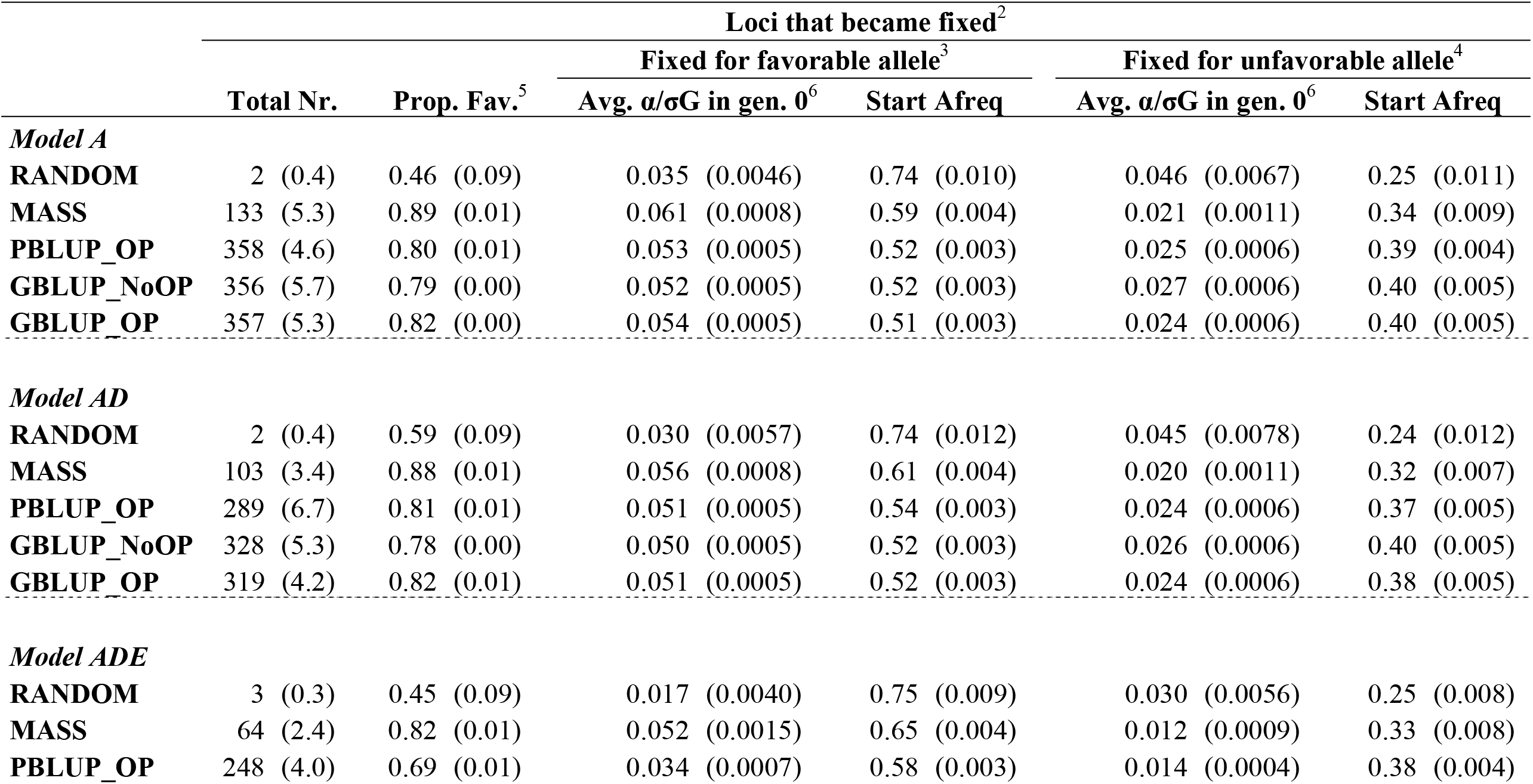
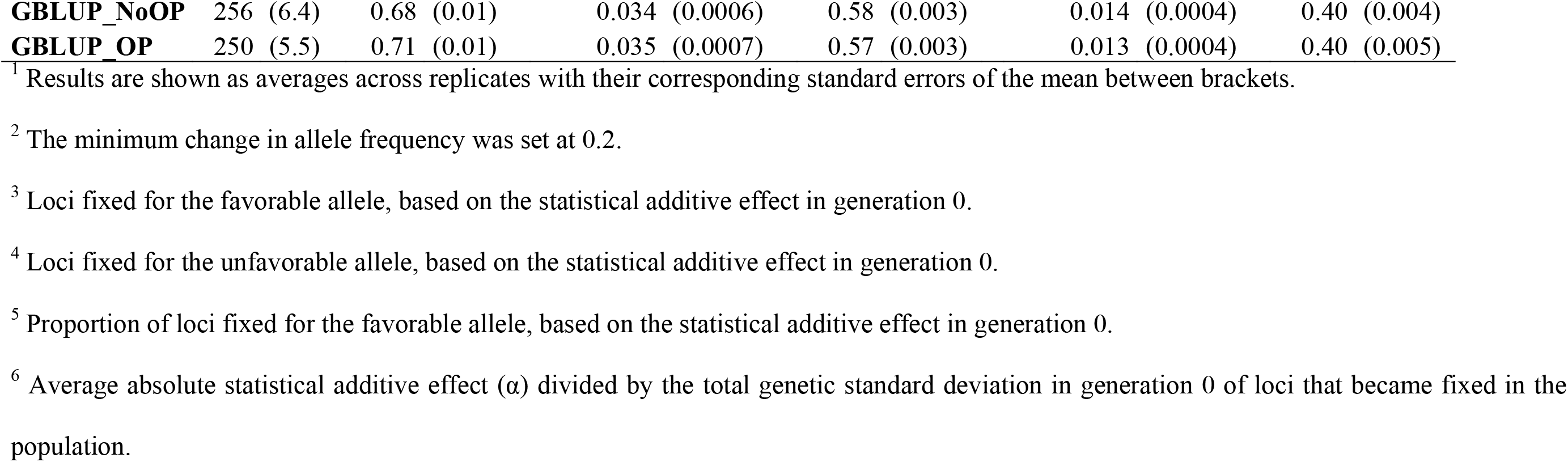
Characteristics of loci that become fixed over 50 generations of selection for the five selection methods and three genetic models^1^. The five selection methods were: RANDOM selection, MASS selection, PBLUP selection with own performance (PBLUP_OP), GBLUP selection without own performance (GBLUP_NoOP) or with own performance (GBLUP_OP). The three genetic models were a model with only additive effects (A), with additive and dominance effects (AD), or with additive, dominance and epistatic effects (ADE).

Besides loci becoming fixed for the favorable allele, with selection a substantial fraction (∼10-30%) of the loci became fixed for the unfavorable allele. The number of loci that became fixed for the unfavorable allele was similar with GBLUP_OP, GBLUP_NoOP and PBLUP_OP, and roughly five times larger than with MASS, while the number of loci that became fixed for the favorable allele was only roughly three times larger with GBLUP_OP, GBLUP_NoOP and PBLUP_OP than with MASS. Moreover, the loci fixed for the unfavorable allele had on average a larger negative statistical additive effect with GBLUP_OP, GBLUP_NoOP and PBLUP_OP than with MASS. For the loci that became fixed for the unfavorable allele, the average frequency of the favorable allele in generation 0 was lowest for RANDOM (∼0.25), followed by MASS (∼0.33), and finally PBLUP_OP, GBLUP_NoOP and GBLUP_OP (∼0.39). This means that, similar to the results for the loci fixed for the favorable allele, the average change in allele frequency at loci where an allele became fixed was largest for PBLUP_OP, GBLUP_NoOP and GBLUP_OP, followed by MASS and finally RANDOM.

### Fixation of loci due to selection versus drift

The average *N_e_*, calculated from pedigree kinship, was largest for RANDOM (222), followed by MASS (143), GBLUP_OP (81), GBLUP_NoOP (72), and lowest for PBLUP_OP (47; Table 5). To investigate the fixation probability due to drift, we simulated populations with those *N_e_* values and without selection, and compared the resulting fixation probabilities to those for the selection scenarios (Figure 7). As expected, with drift only, the highest probability of fixation or loss of the favorable allele was achieved with the lowest *N_e_* (PBLUP), and the lowest probability with the highest *N_e_* (MASS). With drift, the probability of fixation or loss depended only on the starting allele frequency and was independent of its effect. When comparing the fixation probability of selection with drift for a population with the same *N_e_*, we can see that selection increased the probability of a favorable allele to become fixed, especially at intermediate and higher starting frequencies. Moreover, selection resulted in a lower probability of loci to remain segregating.

**FIGURE 7.**
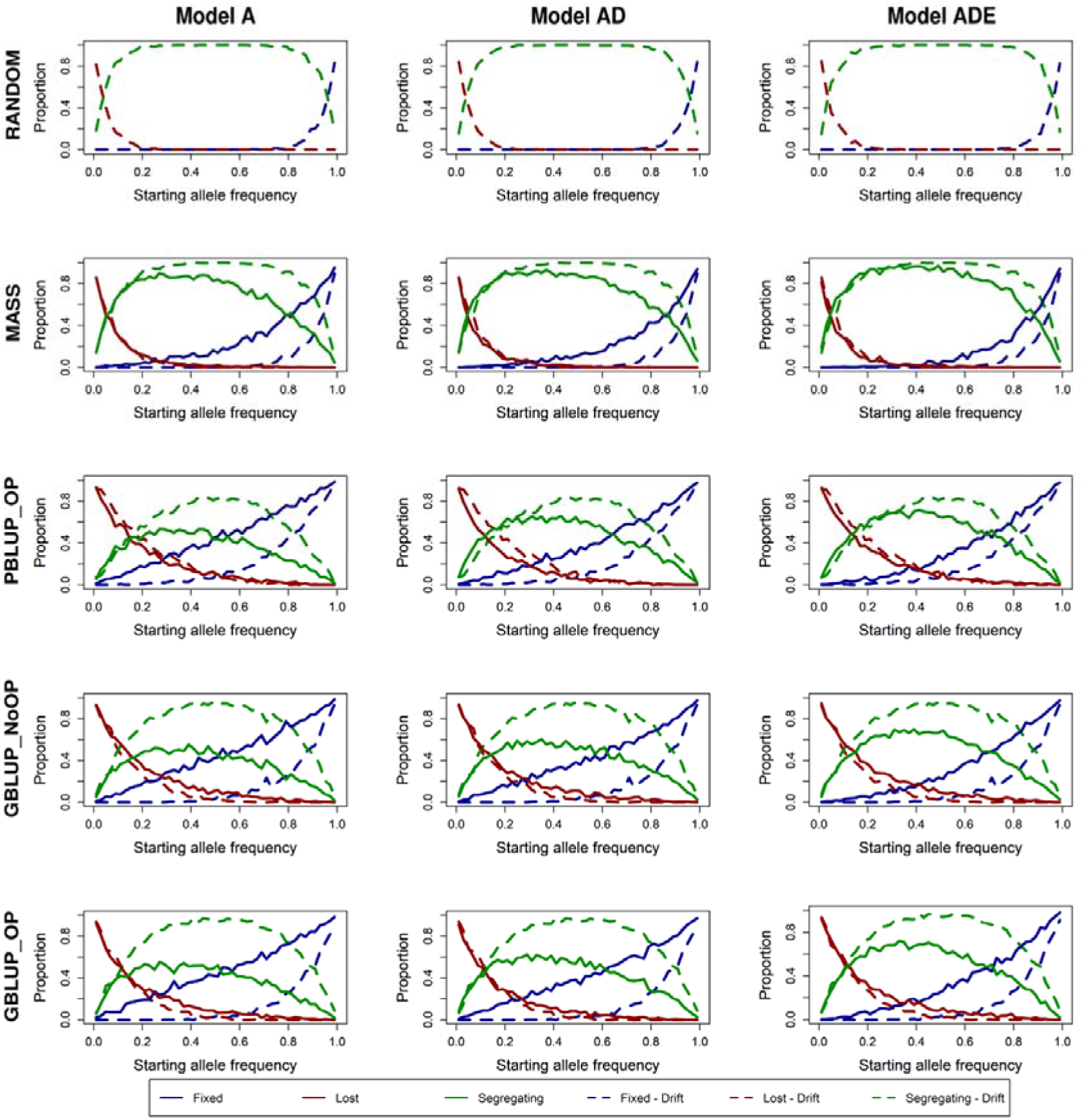
Probability of a locus to become fixed, lost or remain segregating after 50 generations of selection versus 50 generations of drift for the five selection methods and three genetic models. A locus was set to be fixed when the favorable allele was fixed and lost when the favorable allele was lost, where the favorable allele was assigned based on the statistical additive effect in generation 0. The five selection methods were: RANDOM selection, MASS selection, PBLUP selection with own performance (PBLUP_OP), GBLUP selection without own performance (GBLUP_NoOP) or with own performance (GBLUP_OP). The three genetic models were a model with only additive effects (A), with additive and dominance effects (AD), or with additive, dominance and epistatic effects (ADE). Results are shown as averages of 20 replicates.

**TABLE 5.**
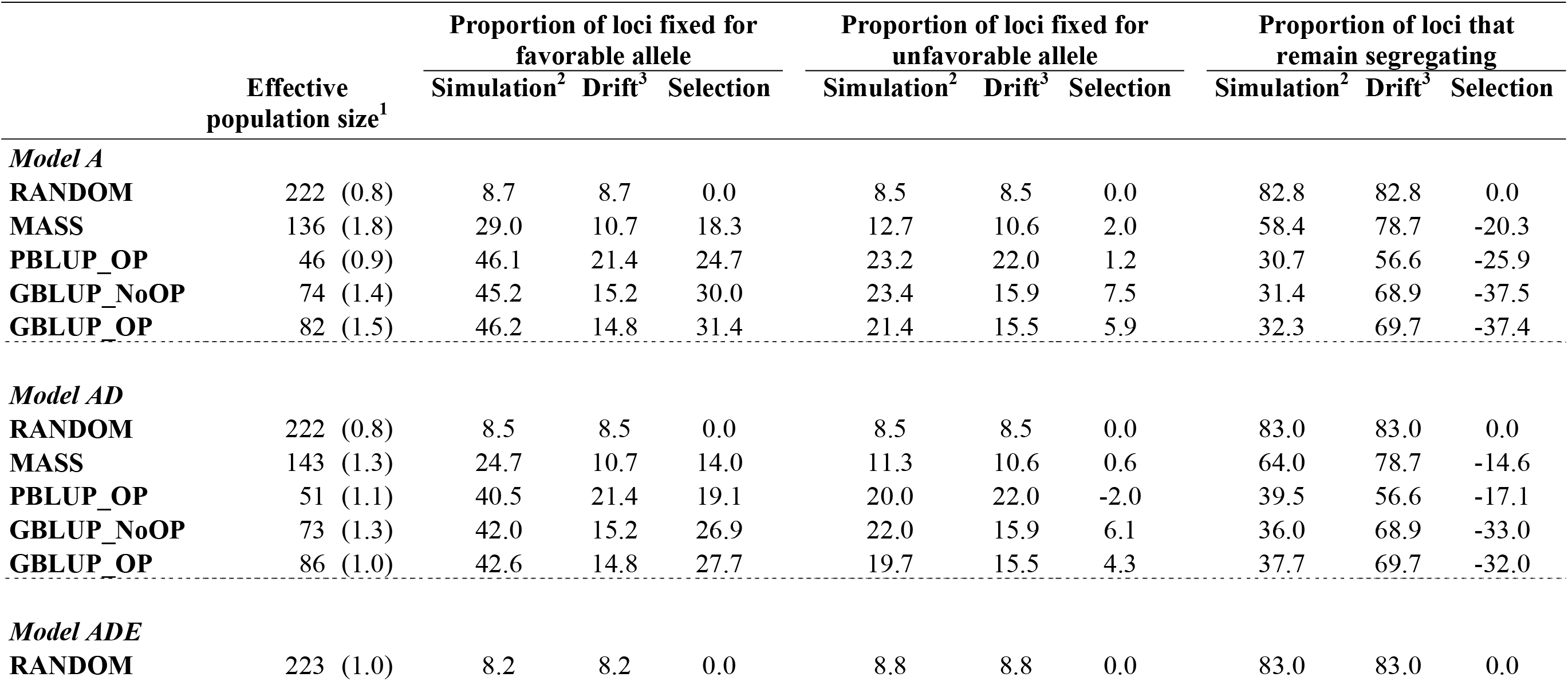
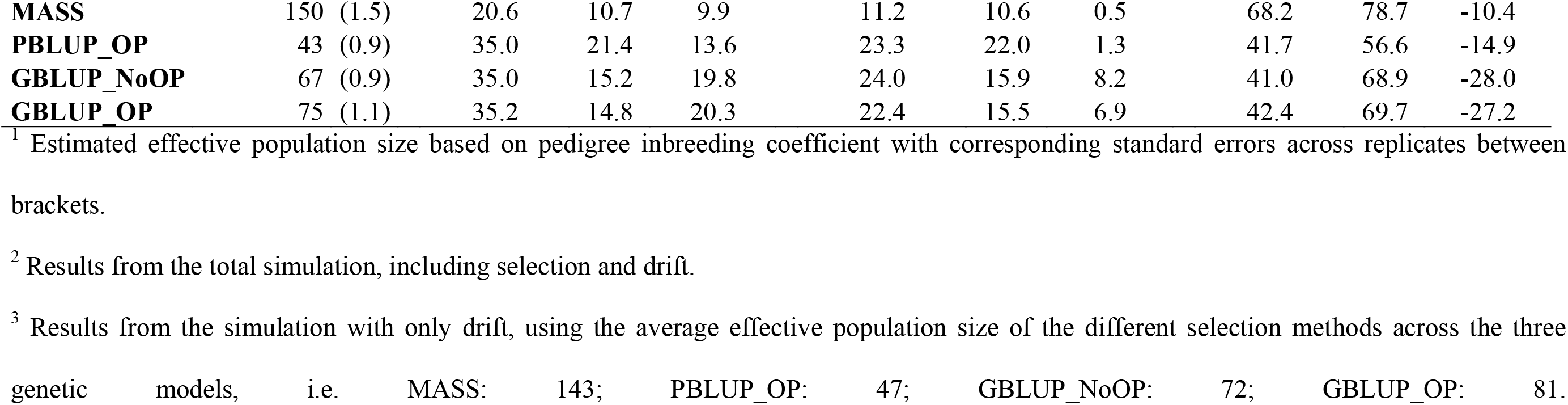
Proportion of loci becoming fixed for the favorable and unfavorable allele or remain segregating as a result of selection and drift for the five selection methods and three genetic models^1^. The five selection methods were: RANDOM selection, MASS selection, PBLUP selection with own performance (PBLUP_OP), GBLUP selection without own performance (GBLUP_NoOP) or with own performance (GBLUP_OP). The three genetic models were a model with only additive effects (A), with additive and dominance effects (AD), or with additive, dominance epistatic effects (ADE).

A quantitative analysis shows that, in general, more loci became fixed for the favorable allele due to selection than due to drift (Table 5). Under MASS, an additional ∼14% of the loci segregating in generation 0 became fixed due to selection compared to drift alone (Table 5). With PBLUP and GBLUP, this proportion was larger, namely ∼19% and ∼26% respectively. Non-additive effects reduced the impact of selection on the fixation of loci. The proportion of the loci segregating in generation 0 that became fixed for the unfavorable allele was approximately equal under drift as under selection for PBLUP_OP. With MASS, an additional ∼1% of the loci lost the favorable allele under selection, while this was ∼6% for GBLUP_OP and ∼7% for GBLUP_NoOP.

### Mutations

The number of mutants that occurred between generations 1 and 49 and that were still segregating in generation 50 was highest with RANDOM, followed by MASS, then by GBLUP_OP and GBLUP_NoOP, and finally by PBLUP_OP (Table 6). For PBLUP_OP, the number of segregating mutants was only ∼45% of that with RANDOM and ∼81% of that with GBLUP_OP and GBLUP_NoOP. The average MAF of the segregating mutants showed the opposite trend (Table 6), with the largest average MAF for PBLUP_OP and the lowest for RANDOM. No clear trend was observed for the variation in MAF of segregating mutants across the different scenarios.

**TABLE 6.**
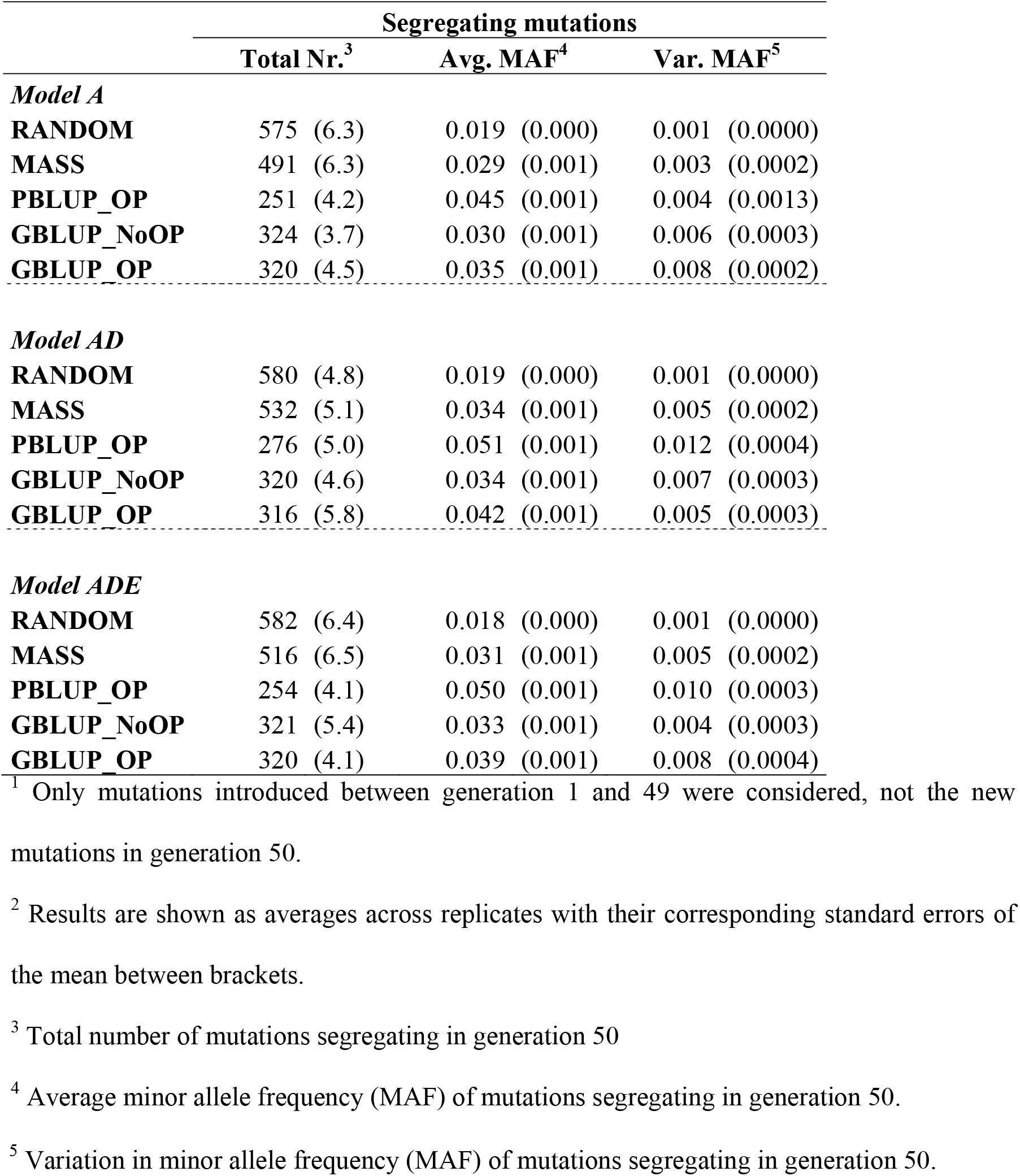
Characteristics of mutations that were segregating in generation 50^1^ for the five selection methods and three genetic models^2^. The five selection methods were: RANDOM selection, MASS selection, PBLUP selection with own performance (PBLUP_OP), GBLUP selection without own performance (GBLUP_NoOP) or with own performance (GBLUP_OP). The three genetic models were a model with only additive effects (A), with additive and dominance effects (AD), or with additive, dominance and epistatic effects (ADE).

|  | Segregating mutations |  |  |
| --- | --- | --- | --- |
|  | Total Nr.<sup>3</sup> | Avg. MAF<sup>4</sup> | Var. MAF<sup>5</sup> |
| *Model A* |  |  |  |
| **RANDOM** | 575 (6.3) | 0.019 (0.000) | 0.001 (0.0000) |
| **MASS** | 491 (6.3) | 0.029 (0.001) | 0.003 (0.0002) |
| **PBLUP_OP** | 251 (4.2) | 0.045 (0.001) | 0.004 (0.0013) |
| **GBLUP_NoOP** | 324 (3.7) | 0.030 (0.001) | 0.006 (0.0003) |
| **GBLUP_OP** | 320 (4.5) | 0.035 (0.001) | 0.008 (0.0002) |
| *Model AD* |  |  |  |
| **RANDOM** | 580 (4.8) | 0.019 (0.000) | 0.001 (0.0000) |
| **MASS** | 532 (5.1) | 0.034 (0.001) | 0.005 (0.0002) |
| **PBLUP_OP** | 276 (5.0) | 0.051 (0.001) | 0.012 (0.0004) |
| **GBLUP_NoOP** | 320 (4.6) | 0.034 (0.001) | 0.007 (0.0003) |
| **GBLUP_OP** | 316 (5.8) | 0.042 (0.001) | 0.005 (0.0003) |
| *Model ADE* |  |  |  |
| **RANDOM** | 582 (6.4) | 0.018 (0.000) | 0.001 (0.0000) |
| **MASS** | 516 (6.5) | 0.031 (0.001) | 0.005 (0.0002) |
| **PBLUP_OP** | 254 (4.1) | 0.050 (0.001) | 0.010 (0.0003) |
| **GBLUP_NoOP** | 321 (5.4) | 0.033 (0.001) | 0.004 (0.0003) |
| **GBLUP_OP** | 320 (4.1) | 0.039 (0.001) | 0.008 (0.0004) |

<sup>1</sup> Only mutations introduced between generation 1 and 49 were considered, not the new mutations in generation 50.
<sup>2</sup> Results are shown as averages across replicates with their corresponding standard errors of the mean between brackets.
<sup>3</sup> Total number of mutations segregating in generation 50
<sup>4</sup> Average minor allele frequency (MAF) of mutations segregating in generation 50.

The cumulative distribution function of the allele frequencies of the mutants shows that more than 90% of the segregating mutants had a frequency below 0.1 across scenarios, except for PBLUP_OP, where this proportion was slightly lower (Figure 8). With RANDOM selection, the maximum allele frequency of a segregating mutant was around 0.4 and no mutants were fixed for the new allele. Over 50 generations of MASS, the maximum allele frequency of a mutant was 0.988. With PBLUP_OP, GBLUP_NoOP and GBLUP_OP, some mutants became fixed (File S2; Table S2.5). The number of mutants becoming fixed was limited for all scenarios, however, ∼40% more mutants became fixed with GBLUP_OP than with PBLUP_OP, and ∼85% less mutants when epistatic effects were present than when only additive effects were present.

**FIGURE 8.**
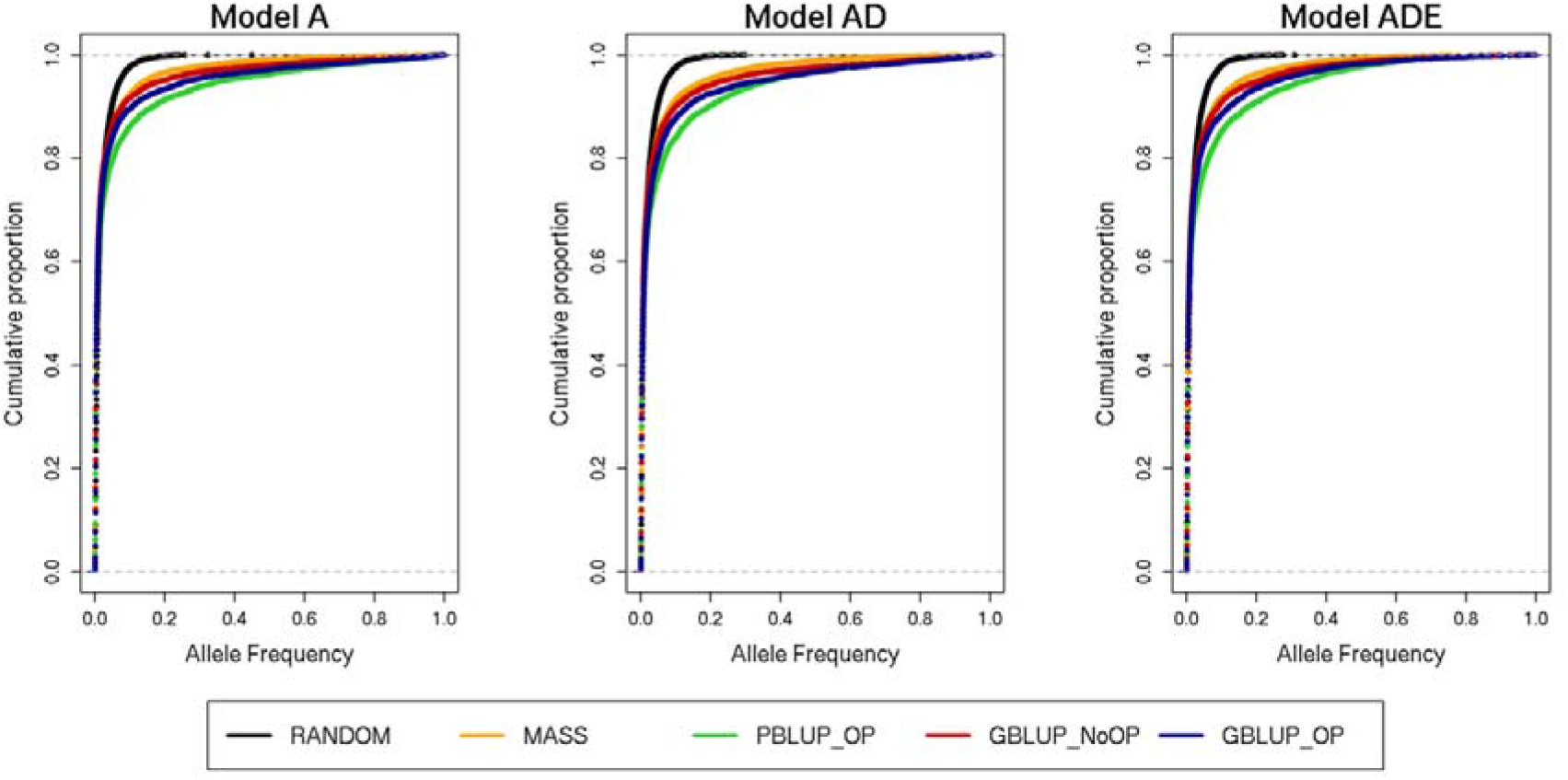
Cumulative distribution function of the allele frequency of the mutated allele in generation 50 of mutations introduced between generation 1 and 49 for the five selection methods and three genetic models. The five selection methods were: RANDOM selection, MASS selection, PBLUP selection with own performance (PBLUP_OP), GBLUP selection without own performance (GBLUP_NoOP) or with own performance (GBLUP_OP). The three genetic models were a model with only additive effects (A), with additive and dominance effects (AD), or with additive, dominance and epistatic effects (ADE). The cumulative distribution function is made across all 20 replicates.

## DISCUSSION

The aim of this study was to investigate the changes in allelic architecture of causal loci and new mutations under 50 generations of random, phenotypic, pedigree, and genomic selection. Moreover, we investigated how those changes were influenced by the genetic architecture of the traits, by investigating traits determined by only additive, additive and dominance, or additive, dominance and epistatic effects. Since the vast majority of causal loci and their interactions are unknown in empirical data, we used simulations. A better understanding of the changes in allelic architecture under selection can help to better predict the long-term effects of selection and may provide clues on how to optimize selection methods to limit undesired loss of genetic variation.

### Changes in allelic architecture

The average change in allele frequency of causal loci under 50 generations of selection was largest with genomic selection (GBLUP), 4% less with pedigree selection (PBLUP), 19% less with phenotypic selection (MASS), and finally 62% less with random selection (RANDOM). This indicates that, as expected, the change in allele frequency was larger when the accuracy of selection was higher. When epistatic effects were present, the average change in allele frequency was 15% lower than for traits only influenced by additive, or additive and dominance effects. In addition, the number of loci with a large change in allele frequency was ∼70% lower when epistasis was present. This is partly a result of the lower narrow-sense heritability for the genetic model with epistasis (∼0.25 for model ADE compared to ∼0.38 for model AD and ∼0.4 for model A), which resulted in a lower selection accuracy. Moreover, epistasis changed the statistical additive effects of loci over generations due to changes in allele frequencies, which also changed the pressure or direction of selection (Falconer and Mackay 1996; Barton *et al*. 2017; Legarra *et al*. 2021; Wientjes *et al*. 2022). Those changes in direction of selection limited the total change in allele frequency. Finally, epistasis resulted in larger average statistical additive effects for loci with a lower MAF (Wientjes *et al*. 2022). This created a negative correlation between statistical additive effects and MAF, which reduced the change in allele frequency at loci with an intermediate frequency. This negative correlation is in agreement with observations in empirical data (Manolio *et al*. 2009; Marouli *et al*. 2017; Zeng *et al*. 2018), and a result of a larger conversion of non-additive effects into statistical additive effects when allele frequencies are closer to 0 or 1 (Barton and Turelli 2004; Hill *et al*. 2008; Mäki-Tanila and Hill 2014). This also means that the additive genetic variance explained by a locus, 2*p*(1 - *p*)*α*^2^, changes due to changes in both *p* and *α* when allele frequency changes (Whitlock *et al*. 1995; Neiman and Linksvayer 2006). We observed that the additive variance was on average highest for loci with a MAF of ∼0.25, and decreased when the MAF of the locus increased further. Thus, non-additive effects can limit the selection pressure on loci at intermediate allele frequencies and can, thereby, limit their change in allele frequency.

Due to the potential changes in selection pressure on loci across generations when epistasis was present (Barton *et al*. 2017), we expected that loci with a large change in allele frequency had on average a lower number of interactions with other loci. Surprisingly, our results showed the opposite and loci with a larger change in allele frequency were generally involved in more interactions (Table 3). This is most likely a result of the positive relation between the number of interactions and the size of the statistical additive effects (File S1; Figure S1.9).

### Effective population size

Selection is known to reduce the effective size of populations (*N_e_*), due to selection of relatives (Robertson 1960; Falconer and Mackay 1996). The lower *N_e_* value for PBLUP indicates that PBLUP resulted in more between-family selection compared to GBLUP and especially MASS, which was also observed before when no measures to restrict inbreeding were taken (Pedersen *et al*. 2010; Liu *et al*. 2014). This is in contradiction to empirical observations, that generally use measures to restrict inbreeding and show an increase in inbreeding per generation since the introduction of genomic selection (Doekes *et al*. 2018; Makanjuola *et al*. 2020; Scott *et al*. 2021). This suggests that the current methods to restrict inbreeding in breeding programs are not working optimally with genomic selection, and that there is a need for new developments (Henryon *et al*. 2019; Meuwissen *et al*. 2020; Woolliams and Meuwissen 2022).

### Fixation of loci

With more accurate selection, i.e., going from MASS to GBLUP, more loci became fixed for the favorable allele. Moreover, selection resulted in an increase in the number of unfavorable alleles becoming fixed, over and above the effect of drift, especially with GBLUP. This was surprising, since selection was expected to prevent favorable alleles from becoming lost compared to drift (Robertson 1960; Walsh and Lynch 2018). This mechanism is probably counterbalanced in the case of MASS and PBLUP and even outperformed in the case of GBLUP by the fixation of unfavorable alleles due to genetic hitchhiking (Smith and Haigh 1974; Barton 2000): the rapid increase in frequency of favorable alleles also changes the frequency of unfavorable alleles at linked loci, because recombination can be too slow to break up the link between loci. As a result, selection can reduce the frequency of a favorable allele when it is linked to unfavorable alleles at other loci (Hospital and Chevalet 1996; Bosse *et al*. 2019). This effect can be considerable and span a large part of the chromosome (Pedersen *et al*. 2010). Therefore, it is desirable but challenging to limit hitchhiking in a breeding program (Sonesson *et al*. 2012).

Our results show that, compared to drift, selection led to fixation of on average an additional 6% (GBLUP_OP) and 7% (GBLUP_NoOP) of unfavorable alleles, which was much larger than with MASS (+ 1%) and PBLUP (+0.1%; Table 5). Those results indicate that the effect of hitchhiking was largest for GBLUP, which is in agreement with previous research showing a stronger hitchhiking effect for GBLUP than for PBLUP (Liu *et al*. 2014). This is probably a balance between different mechanisms. On the one hand, the level of LD among causal loci was slightly stronger for PBLUP than GBLUP (File S1; Figure S1.1), which might result in more hitchhiking for PBLUP. On the other hand, the selection pressure and rate of change of allelic architecture was higher with GBLUP (Figure 5), which might result in more hitchhiking for GBLUP. Moreover, the number of loci becoming fixed for the unfavorable allele due to drift was already much higher for PBLUP, which might limit the additional fixation due to hitchhiking. Overall, the results show that it will be even more important to limit hitchhiking in genomic breeding programs compared to pedigree breeding programs.

With non-additive effects, fewer loci became fixed than under the additive model. Moreover, the proportion of loci that became fixed for the unfavorable allele was higher. At first, we expected that this was because we identified the favorable allele in generation zero, while the favorable allele can change over time (i.e., become the unfavorable allele). However, when we determined the favorable allele based on the average statistical additive effect across the 50 generations, we found very similar results (File S2; Table S2.6). Moreover, while the *proportion* of loci that became fixed for the unfavorable allele was higher with epistasis, the *number* of loci fixed for the unfavorable allele (and also *N_e_*) were similar across genetic models (Table 4). Therefore, the higher *proportion* of loci fixed for the unfavorable allele with epistasis was mainly due to a decrease in the total number of loci that became fixed, which occurred due to a lower selection pressure as a result of a lower narrow-sense heritability.

### Apparent effects

Following our expectations, loci with a larger statistical additive effect in generation 0 showed on average a larger change in allele frequency over 50 generations of selection (Figure 2). Conversely, loci with a large change in allele frequency (>0.9) under selection had on average a larger statistical additive effect (Table 2). However, the change in allele frequency from generation *i* to *i*+1 was only very weakly related to the statistical additive effect in generation *i* (File S1; Figure S1.6). The apparent effect of a locus showed a much higher correlation with the change in allele frequency per generation (Figure 4), indicating that other loci have an impact on the selection pressure on a locus.

The apparent effects as estimated in our study are based on the true breeding values of the individuals. Those breeding values are generally unknown and selection takes place based on estimated breeding values. Therefore, we also estimated the apparent effects based on the estimated breeding values as the *simple* regression of the estimated breeding values on allele counts of a causal locus. The correlation with the change in allele frequency was very similar across selection methods and much larger (∼0.7, File S1; Figure S1.10) than for the apparent effects based on true breeding values (Figure 4). This higher correlation indicates that the impact of selection on the allele frequency at a locus is primarily due to its apparent effect on the EBV, rather than its statistical additive effect. Altogether, those results confirm that selection on a locus is affected by the LD with other causal loci as well as the accuracy of estimating the effect of the locus (Walsh and Lynch 2018).

### New mutations

New mutations will usually be lost by drift within a few generations, largely independent of their effect (Walsh and Lynch 2018). Only when the mutant reaches a certain allele frequency, selection starts to have an effect (Robertson 1960; Lande 1983; Eyre-Walker 2010; Walsh and Lynch 2018). Our results showed that MASS maintained most mutations over 50 generations of selection, followed by GBLUP and finally PBLUP. This is in agreement with previous research, showing that more mutations accumulate in populations with a larger *N_e_* (Hill 1982b; Weber and Diggins 1990). The average MAF of the maintained mutants was highest for PBLUP, followed by GBLUP and finally MASS, which agrees with previous observations for mutations (Mulder *et al*. 2019) and segregating causal loci under selection (Wientjes *et al*. 2022).

MASS resulted in the highest mutational genic variance after 50 generations, followed by GBLUP_OP and PBLUP_OP that showed comparable mutational genic variance, and finally GBLUP_NoOP (File S1; Figure S1.11). This is in agreement with results of Mulder *et al*. (2019), who showed that for maintaining mutational genic variance, own performance records of the selection candidates are important. Interestingly, however, the difference in mutational genic variance between genomic selection with and without own performance was smaller when epistatic effects were present, especially in the last generations. This could be a result of the lower narrow-sense heritability for this genetic model (∼0.25 compared to ∼0.38 for model AD and ∼0.4 for model A (Wientjes *et al*. 2022)), which puts less emphasis on own performance records in the breeding value estimation (Falconer and Mackay 1996).

The mutational genic variance after 50 generations of selection was also compared to the mutational genic variance after 50 generations of drift with the same *N_e_* and an additive model (File S1; Figure S1.12). Results show that the mutational genic variance after 50 generations of drift was very comparable across the different *N_e_* values (range *N_e_* 47-143), because the initial mutational genic variance is independent from *N_e_* (Lande 1976; Hill 1982a). In the first 25 generations of selection, mutational genic variance was similar for all selection methods. Thereafter, MASS increased the mutational genic variance, GBLUP without own performance reduced the mutational genic variance, and PBLUP and GBLUP with own performance maintained the same mutational genic variance compared to drift. Therefore, using own performance information in selection seems to be more important than maintaining a certain *N_e_* to benefit most from new mutations for long-term genetic gain. This benefit, however, became only visible after ∼25 generations of selection. Even though this conclusion might depend on the *N_e_*, it seems to be valid for most livestock populations, since those have similar *N_e_* as the values used here (Hall 2016; Table 1).

## Conclusion

Altogether, our results show that genomic selection results in slightly larger and faster changes in allelic architecture of causal loci than pedigree selection, and much larger and faster changes than phenotypic selection. The presence of non-additive effects limits the change in allele frequency, because non-additive effects can change the selection pressure and direction on a locus over generations. Loci with a larger statistical additive effect changed on average more in allele frequency over 50 generations, however, the correlation between the change in allele frequency and statistical additive effect in one generation was very low for all selection methods. The reason is that the selection pressure on a locus is affected by the linkage phase with other loci and by its allele frequency.

The number of loci that became fixed after 50 generations of selection was comparable for genomic and pedigree selection, but was lower for phenotypic selection. Moreover, compared to phenotypic selection, genomic and pedigree selection had a much higher probability of fixation for unfavorable alleles. With pedigree selection, this increase was almost completely a result of increased drift. With genomic selection, this increase was for a large part due to more hitchhiking, showing the even higher importance to minimize hitchhiking in genomic breeding programs than pedigree breeding programs to limit the loss of favorable alleles that are important for long-term genetic improvement.

Phenotypic selection was best in exploiting new mutations for genetic gain, followed by genomic and pedigree selection with own performance records, and finally genomic selection without own performance records. This shows that own performance records are important to use in selection to optimally exploit new mutations that are essential for long-term genetic gain.

## ACKNOWLEDGMENTS

This publication is part of the project ‘(R)evolution of traits? Quantifying the genetic change in traits over generations as a result of Genomic Selection’ (with project number 16774) of the research programme Veni which is (partly) financed by the Dutch Research Council (NWO). The use of the HPC cluster has been made possible by CAT-AgroFood (Shared Research Facilities Wageningen UR).

